# Attenuation of the 2022 global outbreak monkeypox virus relative to its clade IIb ancestor

**DOI:** 10.1101/2025.02.27.640535

**Authors:** Rebecca P Sumner, Lucy Eke, Bruno Hernáez, Alasdair JM Hood, Telma Sancheira Freitas, Hannah Ashby, Ailish Ellis, Marine Petit, Sian Lant, Isobel Stokes, Preetam Parija, Isabel Alonso, Francisco J. Alvarez-de Miranda, Alazne R. Unanue, Bryan Charleston, Geoffrey L Smith, David O Ulaeto, Antonio Alcamí, Carlos Maluquer de Motes

**Author notes:** Address correspondence to Carlos Maluquer de Motes,.

## Abstract

Monkeypox virus (MPXV) is a zoonotic virus endemic to Africa that has recently re-emerged, becoming the first orthopoxvirus with human-to-human transmission since smallpox eradication. The 2022 global epidemic of MPXV was caused by Clade IIb lineage B.1 virus that derives from the Clade IIb lineage A.1, which is endemic to West Africa^1-3^. Here we compared an early lineage B.1 virus with its closest earlier A.1 ancestor and show that despite only 46 nucleotide differences, lineage B.1 has attenuating phenotypic changes. Both B.1 and A.1 viruses replicated equivalently in cell culture, but B.1 had defects in long-range spread. MPXV B.1 also displayed defects in innate immune control, inducing higher IFNβ, innate immune signalling and MAPK gene expression, and a reduced capacity to suppress interleukin-1β-mediated immune responses. Finally, MPXV B.1 had reduced virulence in a mouse model of MPXV infection relative to the ancestral A.1 endemic strain. Our work demonstrates biological differences between human circulating MPXV lineages that correlate with clinical observations documenting reduced in-host viral dissemination and disease severity for lineage B. Our data also reveal that the APOBEC3-like mutational footprint in MPXV is not only a signature of sustained human transmission but can also drive phenotypic changes.

## MAIN TEXT

MPXV is an orthopoxvirus (OPXV) traditionally confined to West (Clade IIa and IIb) and Central (Clade Ia, Ib) Africa where it infects several rodent species. MPXV causes mpox in humans, a rash and systemic disease similar to smallpox. Historical data report inefficient spread of MPXV and low secondary attack rates even within household contacts^4,5^, supporting the notion that MPXV is poorly adapted to human-human transmission and outbreaks in endemic areas are sustained by periodic re-introductions from the animal reservoir. From May 2022, however, >90,000 cases of mpox were detected outside the African continent, mostly concentrated in the gay, bisexual and men-who-have-sex-with-men (GBMSM) communities, resulting in the declaration of a public health emergency of international concern (PHEIC) by the WHO in July 2022. Little is known about how MPXV (and OPXV in general) emerge and interact with the human host, particularly with regards to innate immunity, which is a critical barrier to zoonotic infection^6-8^. Aside from sporadic infections with cowpox virus, the only human OPXV disease, smallpox (caused by variola virus, VARV) was eradicated >40 years ago and the mechanisms by which it originally emerged are not clear^9^.

Phylogenetic analyses indicate that the Clade IIb MPXV strain that caused the 2022 global outbreak (termed lineage B) is descended from Clade IIb lineage A.1^1-3^ reported from an outbreak occurring in Nigeria between September 2017 and September 2018^10^ and derives from the same zoonotic event^1,3^. Clinical and epidemiological differences exist between lineage B and lineage A strains, with the 2022 global lineage B outbreak being largely sustained by sexual contact transmission and presenting with novel clinical manifestations including a significant primary rash at the infection site, reduced number of lesions in the secondary disseminated rash, and lower case-fatality rate (CFR)^11,12^. Additionally, whilst lineage A viruses have established continuous human-to-human transmission in endemic areas^2^, the prevalence outside West Africa is low and usually associated with defined exportation events^13^. Conversely, lineage B has established an unprecedented global transmission web amongst humans affecting >110 countries. It remains unclear, however, whether these differences are solely explained by non-biological factors or, conversely, distinguishable properties between clade IIb viruses exist.

Lineage B isolates from the first 2022 cases revealed up to 51 unique single-nucleotide changes distributed among them differentiating them from A.1 isolates^1,14^. This equates to a mutation rate of ∼6 changes per year, much higher than expected for a virus estimated to have ∼0.35 nucleotide substitutions per year^15^. Remarkably, these changes are not replication errors, but are consistent with cytosine deaminase activity, likely from host apolipoprotein B mRNA editing enzyme catalytic subunit 3 (APOBEC3) enzymes^3^ and are inherited in the descendant diversity encompassing all known lineage B viruses. More recently, APOBEC3-like mutations have also been reported in a Clade Ib MPXV strain spreading in Central Africa^16,17^. Whilst the APOBEC3-like mutational footprint represents a signature of MPXV transmission amongst humans^3^, it remains unclear whether these mutations impact virus fitness and interactions with the host. Here we took a comparative virology approach to study viral replication and host innate responses to MPXV infection and to determine the biological consequences of the mutational footprint that distinguishes clade IIb lineage A and B MPXV strains.

### Replication and spread of Clade IIb lineage A and B MPXV

In humans, most OPXV infect and/or transmit through direct contact with body fluids and skin lesions, where the virus initially infects keratinocytes and fibroblasts before disseminating within the body through immune cells such as migratory inflammatory monocytes^18,19^. We therefore assessed viral replication following MPXV infection of primary human fibroblasts (HFFF) and THP-1-derived macrophages infected with either Clade IIb, lineage B.1 (isolate CVR_S1) or Clade IIb, lineage A.1 (isolate UK_P3) viruses. OPXV exist in two infectious forms: intracellular mature virus (IMV), a cell-associated form that accumulates inside the cell and is released upon cell lysis, and extracellular enveloped virus (EEV) that are released for dissemination and cell-to-cell spread within the host^19^. Titres of cell-associated virus were comparable between MPXV A.1 and B.1 over 72 h growth curves in primary fibroblasts (Fig. 1a), but the extracellular virus titre upon MPXV B.1 infection was slightly lower, reaching statistical significance at 72 h (Fig. 1b). In THP-1-derived macrophages, cell-associated virus titre was also equivalent, but EEV titres were low for both (Fig. 1c, d). Consistent with HFFF data, MPXV A.1 and B.1 plaque size was equivalent in BSC-40 cells using a semi-solid overlay (Fig. 1e, f), but reduced dissemination by B.1 was observed by ‘comet tail’ assay under liquid overlay, where secondary long-range spread of virus was more visible from MPXV A.1 plaques (Fig. 1g). We also repeated these experiments in the presence of ruxolitinib, an inhibitor of the Janus kinase/signal transducer and activator of transcription (JAK/STAT) pathway, to account for the potential activation of an interferon (IFN) response by B.1 (see below) suppressing virus spread. Ruxolitinib treatment did not rescue EEV titre or comet tail formation by B.1 (Extended Data Fig. 1a, 1c) despite being able to suppress IFN activity in both HFFF and BSC-40 cells (Extended Data Fig. 1b, 1d). Lower extracellular virus titre had been reported for MPXV B.1 relative to Clade I and Clade IIa zoonotic isolates^20^. Our data demonstrates that this is not a general feature of Clade IIb viruses, because Clade IIb lineages A and B viruses differ in this respect.

**Figure 1:**
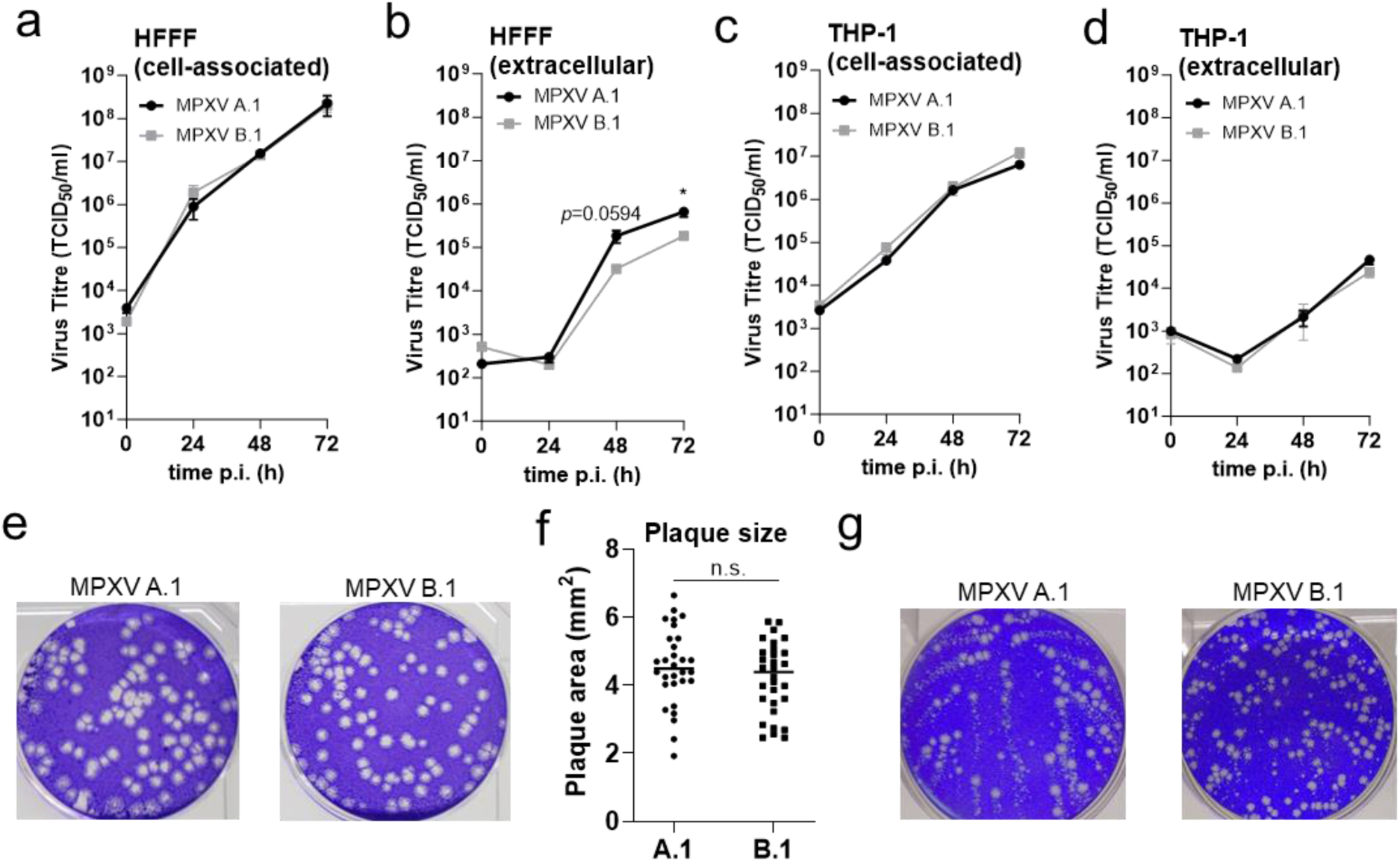
*In vitro* replication of MPXV Clade IIb lineages B.1 and A.1. **a-d**, MPXV A.1 and B.1 replication over 72 h in HFFF (a, b) or THP-1-derived macrophages (c, d) infected at MOI 0.05 quantifying cell-associated (a, c) and extracellular virus (b, d). n = 3 per condition. Data are mean ± SD. Statistical analyses were performed using an unpaired Student’s t-test. \**P* < 0.05. **e**, Representative plaques formed under a semi-solid overlay at 96 hpi on BSC-40 cells. **f**, Plaque area quantification of 30 plaques from e. The horizontal bar represents the median. Statistical analyses were performed using an unpaired Student’s t-test. n.s. non-significant. **g**, Representative comet tail assay from BSC-40 cells infected with MPXV A.1 or B.1 for 96 h with a liquid overlay. All data presented are representative of at least 2 experimental repeats. (g) is representative of 3 repeats.

### Innate immune sensing of MPXV Clade IIb lineages

We next investigated innate immune control by MPXV. Host innate immunity is an important barrier to zoonotic infection, contributes to viral clearance and shapes adaptive immune responses. As cytosolic DNA replicating viruses, poxviruses are susceptible to detection by DNA sensor cyclic 2’3-GMP-AMP (cGAMP) synthetase (cGAS), leading to the production of a second messenger molecule (cGAMP) that binds stimulator of IFN genes (STING) and activates downstream innate transcription factors including IFN responsive factor (IRF)3 to induce IFN expression and subsequent activation of the JAK/STAT pathway^21^. To study host innate responses to the Clade IIb A and B MPXV lineages, viruses were purified by sucrose density ultracentrifugation to minimise carryover of host cell proteins and debris, and experiments were performed at high multiplicity of infection (MOI) to negate differences in viral spread. Both viruses were grown in the same cells at the same time in parallel and tested for protein to infectivity ratios and these were found to be indistinguishable. Hallmarks of cGAS-STING activation including phosphorylation of STING, IRF3 and STAT1 were not observed following MPXV A.1 or B.1 infection in either THP-1-derived macrophages (Fig. 2a) or in primary fibroblasts (Fig. 2b) up to 24 h post-infection, indicating evasion of this pathway. Phosphorylation of these proteins was observed following 2’3’-cGAMP transfection as expected (Fig. 2a, b). Similar results were observed with other OPXV including rodent-restricted ectromelia virus (ECTV) and more promiscuous species like cowpox virus (CPXV) and vaccinia virus (VACV) that are able to infect humans but do not show sustained human-to-human transmission (Fig. 2a, b), in agreement with previous reports^22^. Furthermore, both MPXV strains suppressed STING/IRF3 phosphorylation following exogenous stimulation of infected cells with a dsRNA mimic (poly(I:C)), herring testis (HT)-DNA or 2’3’-cGAMP (Fig 2c), and were equally capable of suppressing HT-DNA-and poly(I:C)-induced *IFNβ* expression by qPCR (Fig 2d). Equivalent infection levels were monitored by blotting for the highly conserved OPXV protein OPG079 (Fig 2a-c). Our data demonstrate that both lineage A and B maintain the ability to inhibit this aspect of host innate immunity.

**Figure 2:**
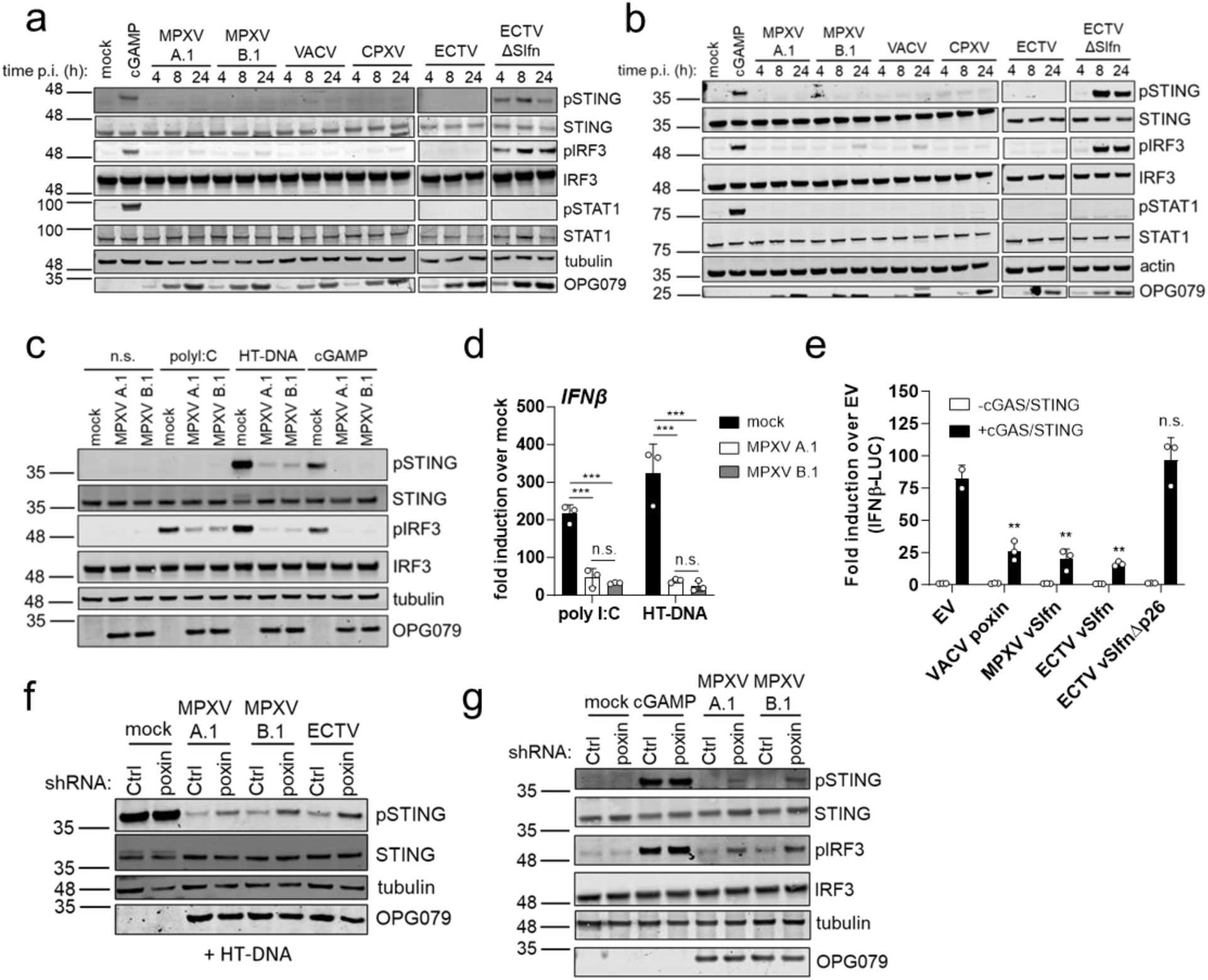
MPXV lineages B.1 and A.1 evade innate sensing in human cells. **a, b** Immunoblot of THP-1-derived macrophages (a) or HFFF (b) mock-infected or infected at MOI 5 for 4, 8 or 24 h with the indicated viruses, or transfected for 4 h with 1 μg/ml 2’3’-cGAMP as a control. **c**, Immunoblot of THP-1-derived macrophages infected for 6 h at MOI 5 with the indicated viruses and then either left non-stimulated (n.s.) or stimulated for 2 h by transfection with 1 μg/ml polyI:C, HT-DNA or cGAMP. **d**, *IFNβ* expression measured by RT-qPCR from HFFF infected with MOI 5 of the indicated viruses for 4 h and then stimulated for 4 h by transfection with 500 ng/ml polyI:C or HT-DNA. Data are mean ± SD. n = 3 per condition. Statistical analyses were performed using an ordinary one-way ANOVA with multiple comparisons. \*\*\**P* < 0.001, n.s. non-significant. **e**, IFNβ reporter activity from HEK293T cells transfected for 24 h with 12.5 ng plasmid expressing VACV poxin, MPXV vSlfn, ECTV vSlfn or ECTV vSlfn lacking the p26 (poxin) domain (ECTV vSlfnΔp26) or an empty vector (EV) control per well and stimulated by co-transfection with 0.6 ng cGAS and 20 ng STING, or 20.6 ng EV as a non-stimulated control. n = 3 per condition. Data are mean ± SD. Statistical analyses were performed using an unpaired Student’s t-test comparing each viral protein to the EV. \*\**P* < 0.01, n.s. non-significant. **f**, Immunoblot of THP-1-derived macrophages stably expressing either a control (Ctrl) or poxin-targeting shRNA, infected for 6 h at MOI 5 with the indicated viruses and then stimulated for 2 h by transfection with 1 μg/ml HT-DNA. **g**, Immunoblot of THP-1-derived macrophages stably expressing either a control (Ctrl) or poxin-targeting shRNA, infected for 8 h at MOI 5 with the indicated viruses or stimulated for 2 h by transfection with 1 μg/ml cGAMP. All data presented are representative of at least 2 experimental repeats. (e) is representative of 3 repeats.

### Viral 2’3’-cGAMP nuclease prevents innate activation by MPXV

OPXV encode a plethora of proteins that antagonise the host immune system, particularly key innate transcription factors such as NF-κB and IRF3, and IFN signalling^23,24^. Most OPXV encode a 2’3’-cGAMP nuclease termed poxvirus immune nuclease (poxin)^25^, which in MPXV, CPXV and ECTV is fused to a C-terminal domain with homology to the human schlafen proteins, hence termed viral Schlafen or vSlfn (Extended Data Fig. 2a, ref. ^26^). Deletion of vSlfn from ECTV was sufficient to induce phosphorylation of STING and IRF3 in infected cells (Fig. 2a, b, ref.^27^) and resulted in severe loss of virulence, concomitant with activation of type I and II IFN responses^27^. The vSlfn protein is conserved (> 90% aa identity) amongst MPXV strains including clade I (Extended Data Fig. 2b). Ectopic expression of MPXV B.1 vSlfn significantly dampened IFNβ activation induced by cGAS/STING to similar levels as ECTV vSlfn and VACV poxin (encoded by OPG188a) (Fig. 2e). ECTV vSlfn lacking the poxin domain (ECTV vSlfnΔp26, Extended Data Fig. 2a) was inactive in this assay as expected^27^ (Fig. 2e). To assess the role of vSlfn during MPXV infection, we used short hairpin (sh)RNA targeting the poxin sequence and validated knock-down against a recombinant VACV expressing FLAG-tagged poxin (Extended Data Fig. 2c). Both MPXV lineages were less able to suppress HT-DNA-induced STING phosphorylation in the absence of vSlfn (shPoxin cells) (Fig. 2f). Furthermore, in contrast to infection in shCtrl cells, MPXV infection in shPoxin cells induced phosphorylation of STING and IRF3 (Fig. 2g), demonstrating that both global and endemic MPXV antagonise IRF3 activation using Poxin.

### MPXV induces a unique inflammatory response that is not observed with other OPXV

The ability of MPXV to cause a disseminated systemic infection and human-to-human transmission is a clear difference from other zoonotic OPXV. To gain a global overview of host responses to MPXV lineages, we performed total RNA sequencing in HFFF and THP-1-derived macrophages infected for 24 h at high MOI. MPXV induced large changes in RNA transcript levels, with around 12-13,000 differentially expressed genes (DEGs, >log2FC 0 and *P*<0.05) with both strains and in both cell types (Fig 3a, b, Extended Data Fig. 3a, b). The top upregulated KEGG pathway in response to MPXV B.1 infection in fibroblasts was MAPK signalling (Fig 3c). Both MPXV strains induced elevated expression of a variety of genes associated with this pathway in HFFF and THP-1-derived macrophages including heat shock proteins (e.g. HSPA6, HSPA7, HSPA1A), dual-specificity phosphatases (e.g. DUSP1, DUSP8) and MAPK transcription factors (Fos, Jun) (Fig 3d). Upregulation of this pathway has previously been reported for other OPXV^24,28^ and in MPXV-infected human skin organoids^29^. In addition, MPXV induced the expression of genes belonging to various cytokine-related pathways including chemokine, TNF and IL-17 signalling (Fig 3c, Extended Data Fig 3c). Infection with either lineage B or lineage A MPXV strains resulted in the elevated expression of a range of inflammatory cytokines belonging to these pathways including ILs 6, 12A and 15 and CXCLs 1-3 and 8 (IL-8), particularly in primary fibroblasts (Fig 3d, e), but also in THP-1-derived macrophages (Fig 3d, f). Concomitantly, significant upregulation of fibrotic cytokine IL-11 was also induced (Fig 3d-f), as was recently observed in MPXV-infected human skin organoids^29^. RT-qPCR confirmed MPXV-induced expression of a subset of these cytokines (*IL6*, *IL11*, *CXCL1* and *CXCL8*) (Fig 3g). Elevated inflammatory markers including TNFα, IL-1β, IL-6 and IL-8, have also been observed in the blood of MPXV-infected individuals^30^. Interestingly, induction of this inflammatory response was unique to MPXV and was not induced, or was poorly induced, by other OPXV including VACV and CPXV, which readily infect humans but rarely establish systemic infection in healthy individuals (Fig. 3g). Although the biological significance of this inflammatory signature is unknown, the clear differential between MPXV and VACV and CPXV identifies it as a potential mechanism for enabling disseminated infection, which itself is a likely enabler of human-to-human transmission. MPXV infection also induced significant downregulation of gene expression and some significantly downregulated pathways were common to fibroblasts and THP-1 cells including cell cycle, DNA replication, base excision repair and lysosome (Fig 3h, i). Notably, enrichment analysis (KEGG) did not identify any significantly upregulated pathways related to IFN or antiviral immune defences for either MPXV strain in either cell type (Fig. 3c, Extended Data Fig 3c-e), in agreement with the significant capacity of these viruses to suppress innate immunity (Fig 2).

**Figure 3:**
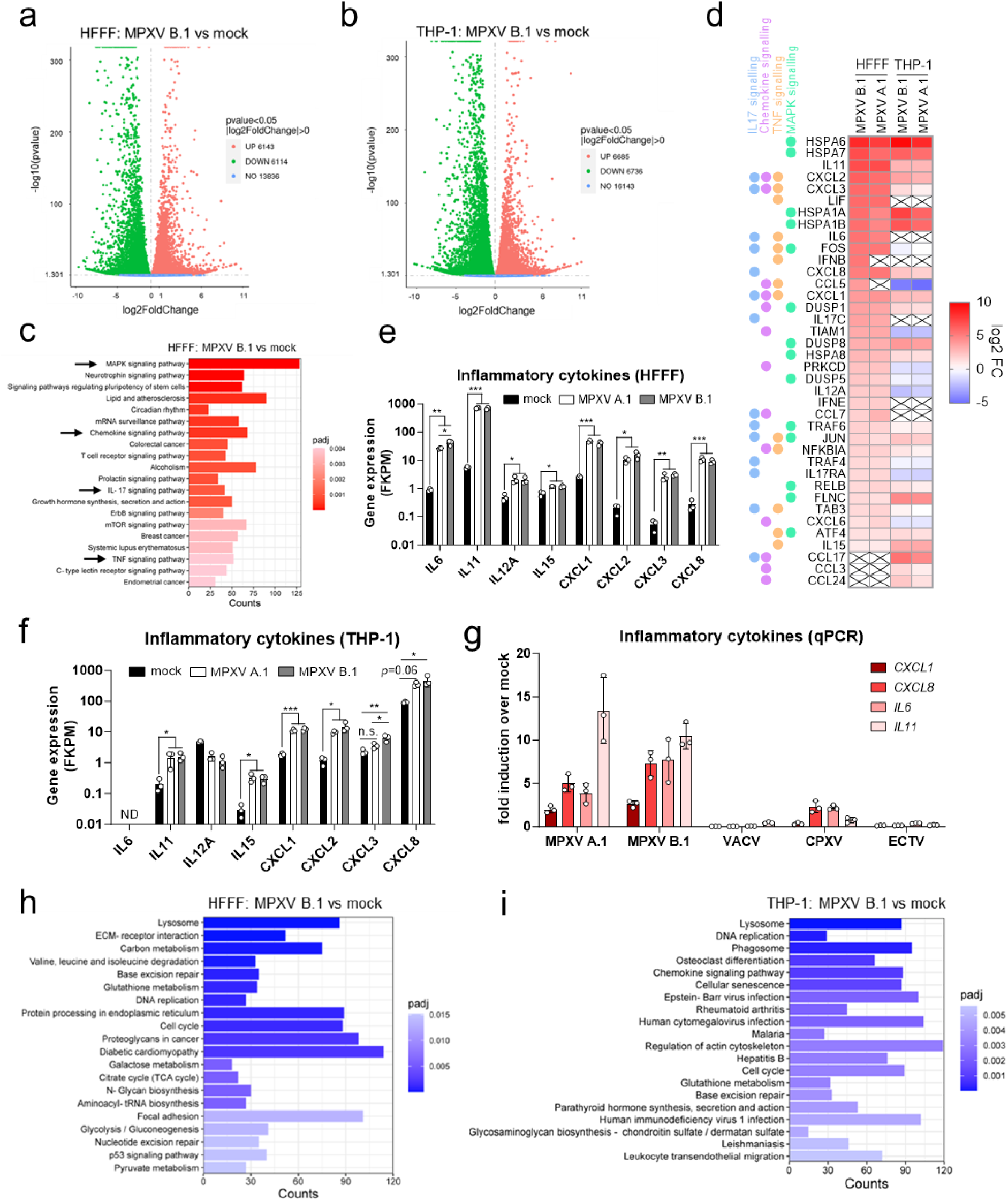
MPXV induces a unique inflammatory response in human cells. **a, b**, Significantly differentially expressed genes (log2FC>0.0) at 24 hpi by volcano plot analysis in HFFF (a) or THP-1-derived macrophages (b) infected with MPXV B.1 at MOI 5, compared with the mock. *P* value determined by DESeq2. **c,** Top 20 significantly upregulated pathways (KEGG analysis) from RNAseq data of HFFF cells infected with MOI 5 MPXV B.1 for 24 h compared with mock-infected cells. *P* value determined by DESeq2. Pathways of interest are highlighted with a black arrow. **d**, Heat map showing log2 fold change (FC) of selected significantly regulated genes from RNAseq data of HFFF or THP-1-derived macrophages infected for 24 h at MOI 5 with MPXV A.1 or MPXV B.1. Boxes with an X indicate transcript not detected. The KEGG pathways to which these genes belong to are indicated on the left. **e, f** Expression level (FKPM) of a panel of inflammatory cytokines identified in **d** in HFFF (e) or THP-1-derived macrophages (f). Data are mean ± SD. n = 3 per condition. Statistical analyses were performed using an ordinary one-way ANOVA with multiple comparisons. \**P* < 0.05, \*\**P* < 0.01, \*\*\**P* < 0.001. **g**, RT-qPCR of selected inflammatory cytokines at 24 hpi from HFFF infected with MOI 5 of the indicated viruses. Data are mean ± SD. n = 3 per condition. **h, i,** Top 20 significantly downregulated pathways (KEGG analysis) from RNAseq data of HFFF (**h**) or THP-1-derived macrophages (**i**) infected with MOI 5 MPXV B.1 for 24 h compared with mock-infected cells. *P* value determined by DESeq2. All data are from a single RNA-seq experiment except (g) which is representative of at least 3 experimental repeats.

### Clade IIb lineage B but not lineage A MPXV induces IFNβ expression in fibroblasts

Although KEGG pathway analysis did not show upregulation of IFN/antiviral defences, MPXV infection did induce the expression of two type I IFNs; β and ε, in primary fibroblasts. Expression of IFNβ, however, was unique to the MPXV lineage B.1 strain and was not induced by the lineage A.1 strain (Fig 3d). A similar result was also observed for the inflammatory chemokine *CCL5*, which was also only induced by MPXV B.1 infection in fibroblasts (Fig 3d). Independent RT-qPCR experiments confirmed induction of these cytokines by MPXV B.1 and no, or very little, induction by MPXV A.1, or indeed other OPXV species, at two MOIs (Fig 4a, b). Type I IFN expression was not however accompanied by global IFN-stimulated gene (ISG) induction in infected cells, and in fact both MPXV lineages greatly suppressed basal ISG expression in both cell types (Fig 4c-e). In addition to blocking ISG induction, viruses are known to counteract IFN restriction by antagonising specific ISGs that are restriction factors^31^. To assess the ability of MPXV to overcome restriction by human ISGs we measured viral replication in HFFFs and THP-1-derived macrophages that had been pre-treated with type I (IFNβ) or type II (IFNγ) IFN to upregulate ISG expression prior to infection. A dose of 5 ng/ml IFNβ (equivalent to ≥50 U/ml) and IFNγ (equivalent to ≥100 U/ml) was found to be sufficient to induce maximal expression of a subset of ISGs (CXCL10, MxA, 2’5’OAS) in both cell types (Extended Data Fig 4). Both MPXV lineages replicated in IFN-treated human cells, reaching viral titres 1-1.5 log lower than in untreated cells (Fig 4f-i).

**Figure 4:**
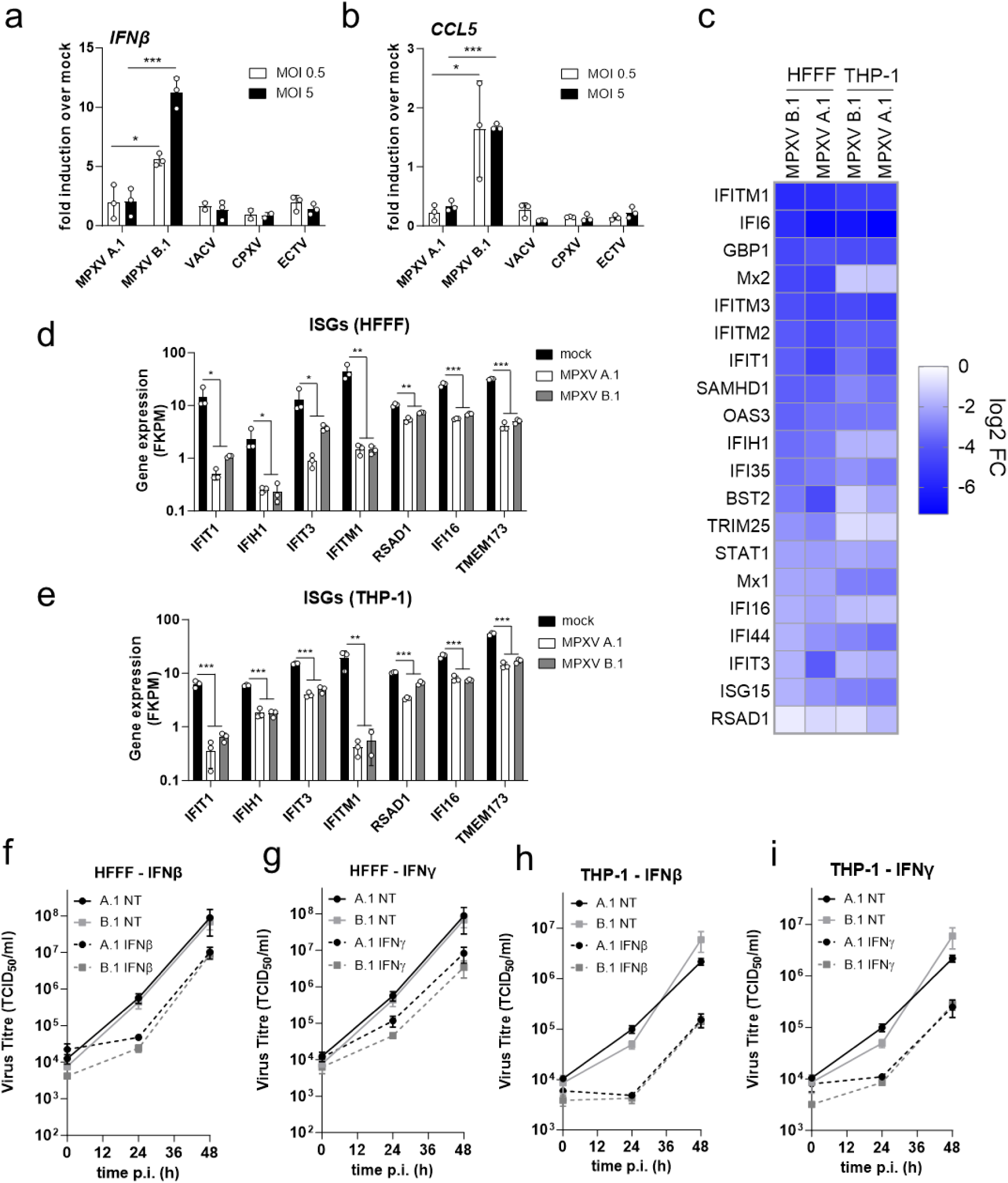
MPXV lineage B.1, but not A.1, induces IFNβ expression in fibroblasts. **a, b,** RT-qPCR for *IFNβ* (**a**) and *CCL5* (**b**) 24 hpi from HFFF infected with MOI 0.5 or 5 of the indicated viruses. Data are mean ± SD. n = 3 per condition. Statistical analyses were performed using an unpaired Student’s t-test. \**P* < 0.05, \*\*\**P* < 0.001. **c,** Heat map showing log2 fold change (FC) of selected ISGs from RNAseq data of HFFF or THP-1-derived macrophages infected for 24 h at MOI 5 with MPXV A.1 or MPXV B.1. **d, e,** Expression level (FKPM) of a panel of ISGs from c in HFFF (d) or THP-1-derived macrophages (d). Data are mean ± SD. n = 3 per condition. Statistical analyses were performed using an ordinary one-way ANOVA with multiple comparisons. \**P* < 0.05, \*\**P* < 0.01, \*\*\**P* < 0.001. **f-i**, Viral titres at the indicated times post-infection from HFFF (g, h) or THP-1-derived macrophages (h, i) pre-treated for 16 h with 5 ng/ml of IFNβ (**f, h**) or IFNγ (**g, i**) or left non-treated (NT) and then infected with the indicated viruses at MOI 0.05. All data are mean ± SD, n = 3 per condition. N.b. NT datasets in f-i are from the same dataset presented in Figs 1a, b. Data presented are representative of 3 (a, b) or 2 (f-i) experimental repeats, except (c-e) that are from a single RNA-seq experiment.

### Differential innate immune control by Clade IIb lineage A and B MPXV

Having observed differences in IFNβ and CCL5 expression in response to infection by lineage A and B MPXV strains, we looked for further differences in host responses to these two viruses. We therefore compared our RNA seq datasets from the two strains and identified a number of DEGs (>log2FC 0 and *P*<0.05) in both cell types (Extended Data Fig 5a, b). Mapping of viral transcripts showed overlapping gene expression patterns for MPXV B.1 and A.1 indicating comparable infection levels (Extended Data Fig. 5c). KEGG enrichment analysis of MPXV B.1 versus A.1 infection identified MAPK signalling as differentially regulated (Fig 5a, b) and numerous genes belonging to this pathway were more upregulated by the B.1 MPXV strain than A.1 in both cell types, indicating differential modulation of this important innate immune pathway by the two strains (Fig 5c, d). These differences were further confirmed by RT-qPCR (Fig 5e). Equally, KEGG pathways differentially regulated by the two strains included various virus infection and innate signalling pathways such as RIG-I-like receptor and TNF signalling (Fig 5a, b). Many of the DEGs within these pathways were innate immune signalling molecules and their expression was more greatly induced by MPXV B.1 than A.1 (Fig 5f). Many of these genes are regulated by the NF-κB family of transcription factors. Poxviruses are well known for their ability to counteract host NF-κB pathways through the expression of a battery of inhibitory proteins^24,32^. We therefore questioned whether the lineage A and B MPXV strains had differential abilities to suppress NF-κB activation and found that whilst both viruses equally modulated TNFα-induced NF-κB-dependent gene expression, the lineage B strain had a reduced capacity to suppress NF-κB-dependent gene expression downstream of IL-1β stimulation (Fig 5g). Equivalent infection levels in these experiments were confirmed by measuring representative early (OPG035), intermediate (OPG057) and late (OPG153) viral transcripts (Extended Data Fig 5d). Taken together, these data reveal differential modulation of host responses by the 2022 lineage B outbreak MPXV relative to its West African Clade IIb ancestor.

**Figure 5:**
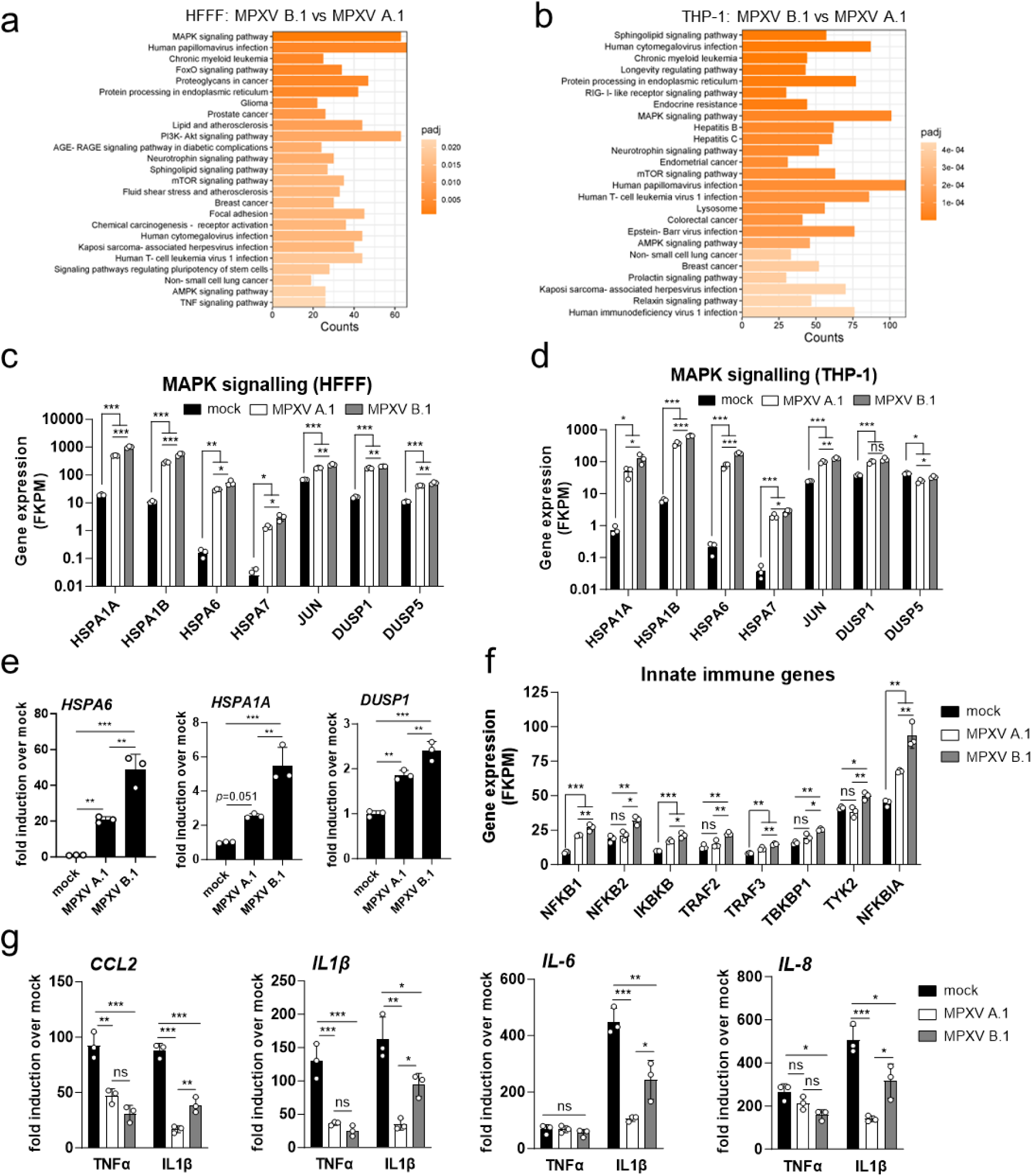
Differential innate immune control by MPXV lineages B.1 and A.1. **a, b,** Top 25 differentially regulated pathways (KEGG analysis) from RNAseq data of HFFF (a) or THP-1-derived macrophages (b) infected with MOI 5 MPXV B.1 for 24 h compared with MPXV A.1-infected cells. *P* value determined by DESeq2. **c, d**, Expression level (FKPM) of a panel of MAPK signalling genes from infected HFFF (c) or THP-1-derived macrophages (d). Data are mean ± SD. n = 3 per condition. Statistical analyses were performed using an ordinary one-way ANOVA with multiple comparisons. \**P* < 0.05, \*\**P* < 0.01, \*\*\**P* < 0.001, ns, non-significant. **e,** RT-qPCR for MAPK signalling genes 24 hpi from THP-1-derived macrophages infected with MOI 5 of the indicated viruses. Data are mean ± SD. n = 3 per condition. Statistical analyses were performed using an ordinary one-way ANOVA with multiple comparisons. \*\**P* < 0.01, \*\*\**P* < 0.001. **f**, Expression level (FKPM) of a panel of innate immune-related genes identified in **c** from infected THP-1-derived macrophages. Data are mean ± SD. n = 3 per condition. Statistical analyses were performed using an ordinary one-way ANOVA with multiple comparisons. \**P* < 0.05, \*\**P* < 0.01, \*\*\**P* < 0.001, ns, non-significant. **g**, RT-qPCR for NF-κB-dependent genes from HFFF cells infected for 4 h with MOI 5 of the indicated viruses and then stimulated for 4 h with either TNFα (50 ng/ml) or IL-1β (10 ng/ml). Data are mean ± SD. n = 3 per condition. Statistical analyses were performed using an ordinary one-way ANOVA with multiple comparisons. \**P* < 0.05, \*\**P* < 0.01, \*\*\**P* < 0.001, ns, non-significant. Data presented are representative of 3 experimental repeats, except (a-d, f) that are from a single RNA-seq experiment.

### Reduced virulence of the 2022 lineage B MPXV in vivo

A reduced ability to disseminate^33^ or to counteract host innate immune responses^27,34^, as observed here for the 2022 lineage B strain relative to its lineage A.1 ancestor, is associated with viral attenuation in small rodent infection models for other OPXV. To assess virulence of these lineages, we infected groups of male and female CAST mice intranasally (i.n.) with either 10^6^ (Fig 6a-c) or 2.5x10^6^ (Fig 6d-f) plaque-forming units (p.f.u.) of each MPXV lineage and monitored animals daily for survival and weight loss. The virus preparations used were grown in the same cell type in parallel, semi-purified in the same way and shown to have the same protein to infectivity ratios, so that any differences seen were likely due to biological differences in the viruses rather than differences in the purity of virus preparations. Under these conditions, infection with lineage A.1 MPXV induced more severe disease than lineage B.1. On average at the higher viral dose, mice infected with A.1 lost 23 % of their body weight 7 days after infection (Fig 6d, e), reaching experimental human endpoints and resulting in a 54 % mortality rate (6/11) (Fig 6f). On the contrary, lineage B.1 MPXV induced 7 % weight loss (Fig 6d, e) and failed to cause mortality (Fig 6f). Furthermore, both lineages exhibited similar replication at the inoculation site (nasal turbinates, Fig 6g). However, viral titres in the distant organs such as the olfactory bulb were significantly lower in MPXV B.1-infected mice indicating a defect in viral dissemination (Fig 6g). Attenuation of the clade IIb MPXV lineage B relative to zoonotic clade IIa and clade I viruses has previously been shown in CAST mice^20^. Our data now reveal virulence differences within clade IIb MPXV, where lineage B.1 is attenuated relative to a closely related lineage A.1 isolate.

**Figure 6:**
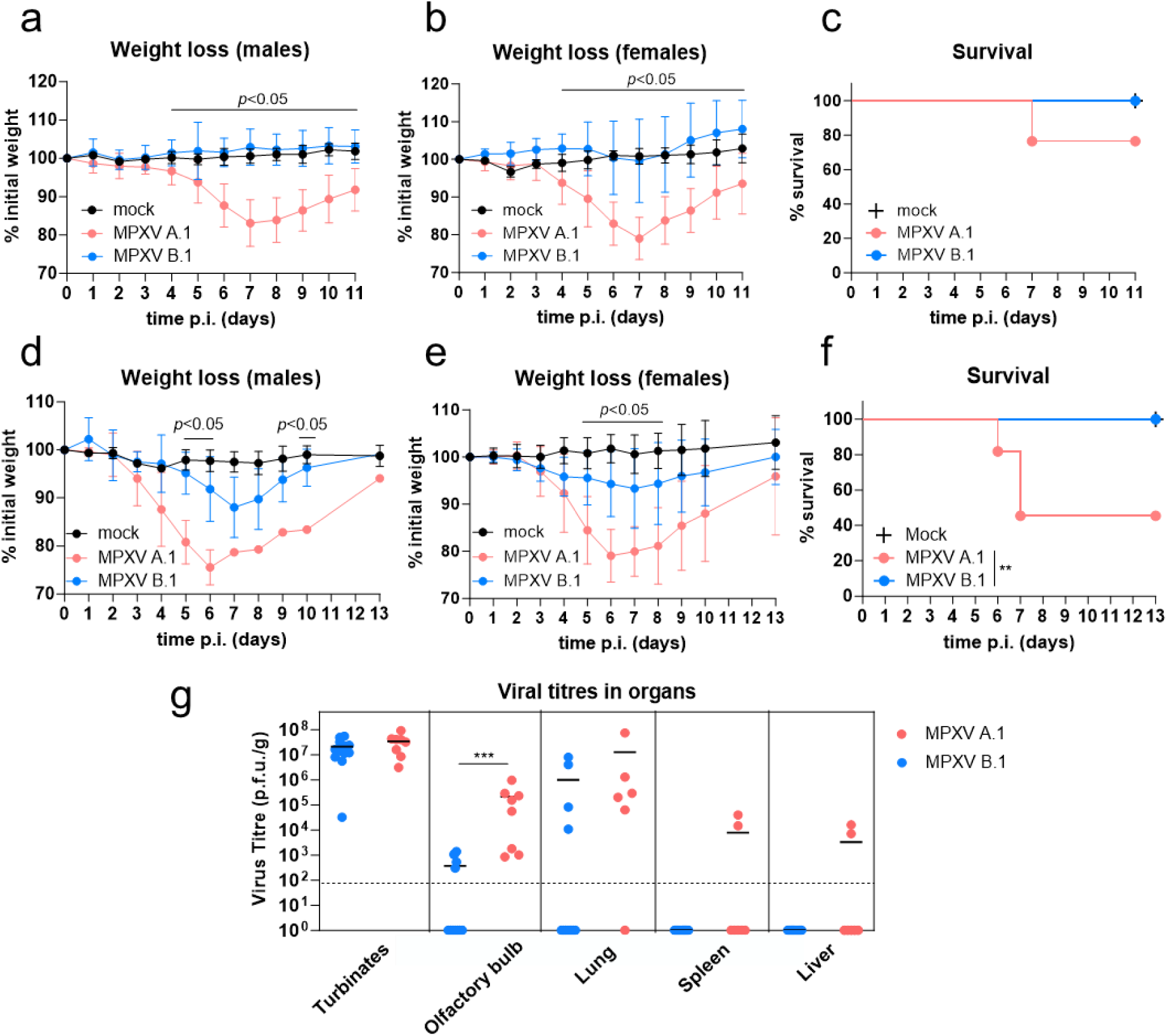
Reduced virulence of MPXV lineage B.1 relative to A.1 *in vivo*. **a, b,** Groups of male (**a,** n=3 mock, n=7 MPXV A.1, n=6 MPXV B.1) and female (**b,** n=3 mock, n=10 MPXV A.1, n=8 MPXV B.1) CAST mice were infected intranasally with 10^6^ p.f.u. of the indicated viruses. Mice were monitored daily for bodyweight and survival. Weight data are expressed as the mean ±SD of the animals’ weights compared to their original weight at the day of inoculation. Statistical analyses were performed using multiple *t*-tests and days at which significant differences (*p* < 0.05) were found between MPXV A.1 and MPXV B.1 are indicated with the solid line. **c** Survival data of males and females combined from **a**, **b**. **d, e,** Groups of male (**d,** (n=3 mock, n=4 MPXV A.1, n=5 MPXV B.1)) and female (**e,** n=4 mock, n=7 MPXV A.1, n=8 MPXV B.1) CAST mice were infected intranasally with 2.5x10^6^ p.f.u. of the indicated viruses and monitored as above. Weight data are presented and analysed (multiple *t*-tests) also as above. **f** Survival data of males and females combined from **a**, **b**. \*\**p* < 0.01 (Mantel-Cox test) **g** Viral titres in organs from CAST mice (n=8 MPXV A.1, n=12 MPXV B.1) 5 days post intranasal infection with 2.5x10^6^ p.f.u. of the indicated viruses. Solid horizontal lines represent the mean titre. Dotted horizontal line indicates limit of detection. *** *p* < 0.001 (Mann-Whitney test). Data represented are from *in vivo* experiments each performed once.

## Discussion

Between September 2017 and September 2018, an outbreak of MPXV clade IIb (now termed lineage A.1) involving 122 confirmed or probable cases was declared in Nigeria^10^. MPXV genomes recovered from Nigeria between the time of the Nigerian outbreak and the 2022 global outbreak revealed an unusual diversification characterised by the accumulation of mutations consistent with APOBEC3 activity^2^. Lineage A has established sustained human-to-human transmission and at present, descendant sub-lineages co-circulate endemically^35^. Although its origin remains obscure, the B lineage descends from the same zoonotic event that caused the Nigerian outbreak and can be phylogenetically traced back to lineage A.1^1-3^. The APOBEC3 family has expanded and diverged into 7 paralogues in humans^36^, whereas only one gene exists in rodents, the major OPXV reservoir. Hence the elevated deamination resulting in the lineage B.1 mutational footprint is consistent with viral exposure to the more complex APOBEC3 system in humans relative to rodents. Certainly, 23/24 amino acid substitutions in 2022 global B.1 MPXV relative to ancestral lineages from 2018 reported here are compatible with APOBEC3 deamination (Supplementary Table 1). At present the distribution of these mutations in MPXV genomes suggest they are random and their presence serves as a marker of human-to-human transmission^2,3^. Our data indicates that, in addition, the mutational footprint in the lineage B MPXV results in both differential spread and host responses, including increased expression of innate immune-related genes such as IFNβ, diminished capacity to counteract host NF-κB responses, and consistent with this, reduced virulence in mice relative to lineage A.1 MPXV. Hence, despite the large 186,000 bp MPXV genome, APOBEC3-like activity has the capacity to drive non-neutral changes in MPXV that result in novel phenotypes. Which of these mutations contribute (individually or collectively) to these differences is the focus of ongoing research. It should be noted though that a number of coding changes are located in genes involved in EEV egress (e.g. OPG056^37^, OPG057^38^) or with known NF-κB and/or MAPK modulatory activity (e.g. OPG047^39^, OPG176^40^), or are located in the termini of the genome which is rich in immune suppressive proteins (Supplementary Table 1).

Our data show that infection with the globally circulating MPXV lineage B.1 results in IFNβ induction and diminished capacity to suppress host innate responses relative to lineage A.1, which so far has not achieved global circulation. Induction of IFNβ did not result in ISG upregulation in infected cells presumably due to the plethora of immune antagonists encoded by OPXV, including the action of a soluble IFN decoy receptor or inhibitors of the IFN signalling pathway, but would be expected to modulate the immune response *in vivo* through paracrine activity on uninfected cells and tissues, establishing an antiviral state. In addition, we observed reduced capacity for B.1 to disseminate *in vitro*. These host responses are consistent with the attenuation in virulence we observed with lineage B.1 in CAST mice and raise questions about their impact on MPXV pathology. For instance, clinical data report 60 % MPXV-infected patients in Nigeria (where lineage A dominates) exhibited substantial viral dissemination with >100 lesions^41^, whereas > 64 % of patients infected with the derivative lineage B.1 MPXV showed fewer than 10 lesions, all localised to the anogenital area^12,42^. Higher lesion count is known to correlate with mpox disease severity^43^. Such reduced dissemination and disease severity in lineage B.1 MPXV is consistent with both reduced extracellular virus and poorer innate immune control. Interestingly, a process of attenuation by loss-of-function is thought to have occurred after the intentional release and subsequent evolution of myxoma virus (a poxvirus of leporids) in Australia in the 1950s to control the introduced European rabbits^44^. Most of these now dominant myxoma virus strains are attenuated and unable to counteract European rabbit PKR due to a single amino acid substitution in one of the viral PKR antagonists^45^. It is thought that loss of immunosuppressive capacity by gene loss also occurred during VARV evolution^46^. Our data suggest a process of attenuation may have happened with Clade IIb MPXV in humans. How this has impacted transmission and global dissemination of mpox, however, remains to be determined and other, non-biological factors are likely to have contributed^47,48^. It is important to note for example that EEV are important for long range spread within the host, but not for host-to-host transmission, which is mediated by IMV^49^. The absence of a generalised disseminated infection as is observed in MPXV B.1-infected patients is expected to influence disease severity, but may have limited impact on transmission if the virus transmits largely via a primary genital rash following sexual contact, and not via the secondary disseminated rashes that follow primary infection in classical MPXV presentations^48^.

The cessation of vaccination against smallpox has resulted in a progressive decline in anti-OPXV herd immunity, and the increase in MPXV zoonoses in endemic areas is a presumed consequence^50^. As anti-OPXV immunity wanes, new outbreaks of poxviruses including MPXV are likely to occur. Understanding the mechanisms that control the evolutionary rates and mediate emergence and adaptation of OPXV to humans is therefore critical. It has been argued that the epidemiology of mpox transmitted via skin primary rash, as seen since 2017, leads to compressed transmission chains relative to historical transmission routes by respiratory droplet or fomites from secondary disseminated rash^48^. In a global setting this could lead to transmission chains being further compressed relative to endemic settings, generating mutations in lineage B.1 faster than lineage A and suggesting this is an effect rather than a cause of global transmission. Consequently, the constant increase of APOBEC3-like mutations is not itself evidence of MPXV adaption to humans. However, this mutational drift provides opportunities for selection, and the data we present here demonstrate differential host responses and virulence between a lineage B.1 virus and a lineage A.1 virus, where 23/24 coding differences are consistent with APOBEC3-like mutations in the lineage B.1 virus. The rate at which deaminase activity edits the MPXV genome appears relatively inefficient^51^. This low level of editing may fail to reach viral error catastrophe in the large MPXV genome as it does in retroviruses, and hence may instead favour the appearance of novel variants. The lineage B.1 viruses used in this study were collected in the UK and Spain in June 2022 shortly after the emergence of MPXV in non-endemic countries in May 2022, suggesting that this lineage had acquired the attenuated phenotypes reported here prior to, or early in, the global outbreak, and descendant lineages are likely to have them as well. It is conceivable that milder disease is less likely to limit host mobility or require major medical attention prompting spread and underdetection^47^. The ability of host enzymes to generate novel phenotypic profiles is therefore an important consideration for surveillance and control of current and future MPXV epidemics. Our observations are underscored by the recent emergence of a novel MPXV clade I variant that is also rich in APOBEC3-like mutations that may contribute to different virulence and transmission rate^16,17^.

## METHODS

### Cells and reagents

BSC-40, HFFF2 and HEK293T cells were cultured in Dulbecco modified Eagle medium (Sigma) and THP-1-IFIT1-Gluc cells (modified to express Gaussia luciferase under the control of the endogenous *IFIT1* promoter^52^) were cultured in RPMI medium (Gibco), all supplemented with 10 % foetal bovine serum (FBS, Biowest) and 100 U/mL penicillin and 100 µg/mL streptomycin (Pen/Strep; Life Technologies). THP-1 cells were differentiated for 48 h in the presence of 50 ng/ml PMA (Peprotech). HT-DNA (Sigma), 2’3’-cGAMP (Invivogen) and polyI:C (Invivogen) were used to stimulate cells by lipofectamine 2000 (Invitrogen) transfection according to the manufacturer’s instructions. Human IFNβ and γ were obtained from Peprotech and dissolved in water at 100 µg/ml (equivalent to ≥10^6^ U/ml for IFNβ and ≥2x10^6^ U/ml for IFN γ). Human TNFα and IL-1β were also from Peprotech. Universal type I IFN was from PBL Assay Science. JAK inhibitor ruxolitinib was purchased from Cambridge Bioscience.

### Virus stocks and cell infections

MPXV Clade IIb lineage B viruses (MPXV_CVR_S1; accession number ON808413 and MPXV_CBM; accession number GCA_964276875) were isolated in 2022 from patients at the University of Glasgow-Centre for Virus Research (CVR) and at the Centro de Biologia Molecular Severo Ochoa (CBM) in Madrid, Spain respectively, and differ in 1 coding change. Lineage A viruses (MPXV_UK_P2; accession number MT903344 and MPXV_UK_P3; accession number MT903345) were isolated in 2018 at the UK Health Security Agency and are genetically identical. Viruses were amplified in BSC-40 (MPXV_CVR_S1 and MPXV_UK_P3) and BSC-1 (MPXV_CBM and MPXV_UK_P2) cells before purification and titration by plaque assay. Both MPXV_CVR_S1 and MPXV_CBM isolates were characterised for viral growth with indistinguishable results between them and relative to their MPXV_UK_P2 and MPXV_UK_P3 lineage A comparators. Other viruses used were VACV Western Reserve (AY243312), CPXV Brighton-Red (AF482758.2), and ECTV Moscow (AF012825). ECTVΔvSlfn has been described^27^. VACV, CPXV and ECTV were expanded in BSC-40 cells and purified before titration by plaque assay as described previously^53^. Purification was performed by ultracentrifugation of cytoplasmic extracts over a 36 % (w/v) sucrose cushion (32,900 × g for 80 mins at 4 °C) prior to use. Protein-to-infectivity ratios were calculated for all MPXV isolates and shown to be indistinguishable. All infections were performed in medium supplemented with 2.5 % FBS. Cells were initially infected in a minimal volume of medium on ice for 60-90 mins before unbound virus was washed off with medium and cells transferred to 37 °C. All infection experiments were performed in a Biosafety level 3 laboratory at The Pirbright Institute, Surrey. Plaque size was measured using ImageJ.

### IFN sensitivity assays

For IFN sensitivity assays cells were seeded at 2.5x10^5^ cells/well in 24 well plates. The following day cells were treated overnight with 5 ng/ml IFNβ or γ, or mock treated with medium and then infected the next morning on ice at an MOI 0.05 in 200 μl/well medium. After 60 min the inoculum was removed and the cells washed once with medium to remove unbound virus before 500 μl medium was added per well and the cells transferred to 37 °C. Virus was harvested at 0, 24 and 48 hpi by scraping the cells in their medium and freeze-thawing three times before calculation of viral titres by tissue culture infectious dose 50 (TCID50) assay according to the Reed and Muench method as previously described^54^. TCID50 assays were performed on BSC-40 cells and incubated for 6-7 days.

### RT-qPCR

For qPCR cells were seeded at 2.5x10^5^ cells/well in 24 well plates and harvested in 140 μl PBS + 560 μl buffer AVL (QIAgen) containing 1 % (v/v) triton X-100. After incubation at room temperature for 10 mins, 560 μl ethanol was added and incubated for a further 20 mins before extracting total RNA using the RNeasy mini RNA extraction kit (QIAgen) including on-column DNase treatment according to the manufacturer’s instructions. Five hundred ng RNA was used to synthesise cDNA using Superscript III reverse transcriptase (Invitrogen), also according to the manufacturer’s protocol. cDNA was diluted 1:5 in water and 1.5 μl was used as a template for real-time PCR in a 5 μl reaction volume total containing SYBR® Green PCR master mix (Applied Biosystems). Gene expression was quantified on a Quant Studio 5 real-time PCR machine (Applied Biosystems). Expression of each gene was normalised to an internal control (*18S RNA*) and these values were then normalised to mock-treated control cells to yield a fold induction. The following primers were used to detect host genes:

18S RNA: Fwd 5’-GTAACCCGTTGAACCCCA-3’, Rev 5’-CCATCCAATCGGTAGTAGCG-3’ CXCL-10: Fwd 5’-TGGCATTCAAGGAGTACCTC-3’, Rev 5’-TTGTAGCAATGATCTCAACACG-3’ MxA: Fwd 5’-CCCCAGTAATGTGGACATCG-3’, Rev 5’-ACCTTGTCTTCAGTTCCTTTGT-3’ 2’5’ OAS: Fwd 5’-TGTGTGTGTCCAAGGTGGTA-3’, Rev 5’-TGATCCTGAAAAGTGGTGAGAG-3’ CXCL1: Fwd 5’-CTGCGCTGCCAGTGCTTGCA-3’, Rev 5’-TGTGGCTATGACTTCGGTTTG-3’ CXCL8 (IL-8): Fwd 5’-AGAAACCACCGGAAGGAACCATCT-3’, Rev 5’-AGAGCTGCAGAAATCAGGAGGGCT-3’ IL6: Fwd 5’-AAATTCGGTACATCCTCGACG-3’, Rev 5’-GGAAGGTTCAGGTTGTTTTCT-3’ IL11: Fwd 5’-GGACATGAACTGTGTTTGCC-3’, Rev 5’-CCGTCAGCTGGGAATTTGTC-3’ IL1β: Fwd 5’-ACAGATGAAGTGCTCCTTCCA-3’, Rev 5’-GTCGGAGATTCGTAGCTGGAT-3’ ICAM1: Fwd 5’-TCTGTGTCCCCCTCAAAAGTC-3’, Rev 5’-GGGGTCTCTATGCCCAACAA-3’ IFIT1: Fwd 5’-CCTGAAAGGCCAGAATGAGG-3’, Rev 5’-TCCACCTTGTCCAGGTAAGT-3’ IFNβ: Fwd 5’-ACATCCCTGAGGAGATTAAGCA-3’, Rev 5’-GCCAGGAGGTTCTCAACAATAG-3’ CCL5: Fwd 5’-CCCAGCAGTCGTCTTTGTCA-3’, Rev 5’-TCCCGAACCCATTTCTTCTCT-3’ HSPA6: Fwd 5’-GATGTGTCGGTTCTCTCCATTG-3’, Rev 5’-CTTCCATGAAGTGGTTCACGA-3’ HSPA1A: Fwd 5’-AGCAGGTGTGTAACCCCATC-3’, Rev 5’-GCAGCAAAGTCCTTGAGTCC-3’ DUSP1: Fwd: 5’-GCTCAGCCTTCCCCTGAGTA-3’, Rev 5’-GATACGCACTGCCCAGGTACA-3’ CCL2: Fwd 5’-CAGCCAGATGCAATCAATGCC-3’, Rev 5’-TGGAATCCTGAACCCACTTCT-3’

Primers to detect viral genes were as follows:

OPG035: Fwd 5’-ATCATCCTAAAGAGTATGGTG-3’, Rev 5’-GCTAGATAGATTGAATCAACC-3’ OPG057: Fwd 5’-CGATTTACTGTGGCTAGATAC-3’, Rev 5’-ATATCACTTCGGCAAATTTCG-3’ OPG153: Fwd 5’-ATCTCCCATGTGGTGGAATAC-3’, Rev 5’-GTTGATAGGTTAGAACATCAC-3’

### Transcriptomics and bioinformatics

For transcriptomics experiments cells were seeded at 2.5x10^5^ cells/well in 24 well plates and infected in triplicate with MPXV A.1, MPXV B.1 or mock infected as a control at an MOI 5 for 24 h. RNA was purified as described above using the RNeasy mini RNA extraction kit with on-column DNase treatment and sent for bulk RNA sequencing by Novogene. Messenger RNA was purified from total RNA using poly-T oligo-attached magnetic beads. After fragmentation, the first strand cDNA was synthesised using random hexamer primers, followed by the second strand cDNA synthesis using either dUTP for directional library or dTTP for non-directional library. The library was checked with Qubit and real-time PCR for quantification and bioanalyzer for size distribution detection. Quantified libraries were then pooled and sequenced on an Illumina platform. RNA-seq reads were aligned to the *Homo Sapiens* genome using HISAT2^55^ and featureCounts v1.5.0-p3 was used to count the read numbers mapped to each gene and the FPKM of each gene was calculated based on the length of the gene and reads count mapped to it. Differentially expressed genes (DEG) were calculated using DESeq2^56^. The resulting *P*-values from DESeq2 were adjusted using the Benjamini and Hochberg’s approach for controlling the false discovery rate. Genes with an adjusted *P*-value <0.05 found by DESeq2 were assigned as differentially expressed. DEGs were then used for KEGG pathway enrichment analysis^57^. Pathway *p*-values <0.05 were considered significant. Heat maps were generated using GraphPad Prism (version 10) and KEGG pathway analysis figures were generated using SRPlot (accessed from https://www.bioinformatics.com.cn/srplot). For mapping of viral gene reads, RNA sequencing reads were aligned to a MPXV reference genome (NC_063383.1) using the HISAT2 software. Following alignment, the featureCounts tool was employed to quantify the read counts mapped to each gene. These counts were then normalised to counts per million (CPM) to facilitate comparison across samples.

### shRNA lentivirus transduction

Hairpin sequences against the VACV poxin gene (OPG188a) were designed using siRNA Wizard v3.1 Software (InvivoGen) and annealed into oligo duplexes. The duplexes were then cloned into an HIV-1-based shRNA expression vector encoding puromycin resistance (pSIREN, Clontech) at BamHI-EcoRI sites, and the products were checked by sequencing. VSV-G-pseudotyped lentiviral vectors were produced by co-transfecting HEK293T cells in 10-cm plates with 1.5 μg of the shRNA plasmid, 1 μg p8.91^58^ (encoding Gag-Pol, Tat and Rev), and 1 μg pMDG (Genscript) encoding VSV-G using Fugene6 (Promega) according to the manufacturer’s instructions. The medium was changed the following day and the supernatant harvested at 48 and 72 h post-transfection. THP-1 cells were transduced with neat supernatant by spinoculation (1000 × g for 1 h at room temperature) and a polyclonal cell line selected in the presence of puromycin (1 μg/ml). Poxin shRNA target sequences:

Top: GAAGGAGTAGGGATTCATCAT Bottom: CAAAGAGAAGGCCAAAGAAAT

### Construction of a VACV expressing FLAG-tagged poxin

For construction of VACV encoding Flag-tagged OPG188a (poxin) from its natural locus (VACV.Flag-poxin), a transient dominant selection strategy was used. First, we generated a plasmid containing the coding sequence of *OPG188a* fused to 3 repeats of the Flag-tag at the N terminus, flanked by 200-bp of the 5’ and 3’ regions of the *OPG188a* locus, and the Escherichia coli guanylphosphoribosyl transferase (Ecogpt) gene fused in-frame with the enhanced green fluorescent protein (EGFP) gene (pFlag-poxin)^34^. BSC-40 were infected with VACV strain Western Reserve lacking *OPG188a* ^59^ and then transfected with pFlag-poxin. Enhanced green fluorescent protein (EGFP)-positive plaques were then selected in the presence of mycophenolic acid, hypoxanthine, and xanthine, as described previously^60^. The genotype of the resolved viruses was analysed by PCR following lysis and proteinase K treatment of infected BSC-40 cells using primers that anneal to the flanking regions of *OPG188a*.

### Reporter gene assays

Dual luciferase reporter gene assays in HEK293T cells were performed as we have described previously^27^. Plasmids encoding codon-optimised ECTV vSlfn and ECTV vSlfn lacking the poxin/p26 domain (ECTV vSlfnΔp26) have also been previously described^27^. Codon-optimised VACV *OPG188a* (encoding poxin, ordered from GeneArt, Invitrogen) and MPXV *OPG188* (encoding vSlfn, obtained from Mike Weekes, University of Cambridge) were cloned into a modified pcDNA4 vector with 3 copies of the FLAG tag at the N terminus^61^. Gaussia luciferase (GLuc) activity in supernatants from THP-1-IFIT1-GLuc was measured by transferring 10 μl to a white 96 well assay plate, injecting 50 μl per well of coelenterazine substrate (Nanolight Technologies, 2 μg/ml) and analysing luminescence on a CLARIOstar plate reader (BMG Labtech). Fold inductions were calculated by normalising to a mock-treated control.

### Immunoblotting

For immunoblotting cells were seeded at 5x10^5^ cells/well in 12 well plates and harvested in 120 μl RIPA buffer (50 mM Tris-HCl pH8, 150 mM NaCl, 1 % (v/v) NP-40, 0.5 % (w/v) sodium deoxycholate, 0.1 % (w/v) SDS) supplemented with protease and phosphatase and protease inhibitors (Roche) and benzonase (EMD Millipore). Samples were boiled for 5 mins in boiled in Laemmli buffer (final concentration 50 mM Tris-HCl pH 6.8, 2 % (w/v) SDS, 10 % (v/v) glycerol, 0.1 % (w/v) bromophenol blue, 100 mM betamercaptoethanol) and proteins were separated by SDS-PAGE on 4-12 % bis-tris polyacrylamide gels (Invitrogen). After PAGE, proteins were transferred to a Hybond ECL membrane (Amersham biosciences) using a semi-dry transfer system (Biorad). Primary antibodies were from the following sources: pSTING Ser366 (Cell Signaling Technologies 19781S), STING (Cell Signaling Technologies 13647S), pIRF3 Ser386 (Abcam AB76493), IRF3 (Abcam AB68481), pSTAT1 Tyr701 (Cell Signaling Technologies 9167S), STAT1 (Cell Signaling Technologies 14994S), IFIT1 (Cell Signaling Technologies 14769S), MxA (Cell Signaling Technologies 37849S), tubulin (EMD Millipore 05-829), actin (Proteintech 66009-1-Ig), FLAG (Sigma F1804), OPG079 (a gift from D. Ulaeto, Defence Science and Technology Laboratory). Primary antibodies were detected with goat-anti-mouse/rabbit IRdye 680/800 infrared dye secondary antibodies and membranes imaged using an Odyssey Infrared Imager (LI-COR Biosciences).

### MPXV infection of mice

The experiments described and CAST/EiJ mice breeding were carried out at the Centro de Biologia Molecular Severo Ochoa (CBM) in Madrid (Spain) and were approved by the Ethical Review Board of CBM and CSIC under reference PROEX 241.1/21. Groups of 7-12 weeks (12-16 g) male and female CAST/EiJ mice were intranasally (i.n.) infected with a single 10 μl dose containing 10^6^ or 2.5x10^6^ p.f.u. of MPXV lineage A.1 (MPXV_UK_P2) or lineage B.1 (MPXV-CBM). After infection, viral inocula used were back-titrated by plaque assay to verify the administered viral dose. Mice were housed in ventilated racks under biological safety level 3 containment facilities and monitored daily for survival, weight, temperature and signs of disease. Determination of virus titres in organs from infected mice was performed at 5 days post-infection. Tissue sample extraction and processing was performed as previously described^62^. Infected mice were euthanised by CO2 asphyxiation, and a sample of the indicated organs from each animal were aseptically removed. Samples were weighed, homogenised in 1 ml PBS, and freeze/thawed 3 times. Then, the amount of infectious virus was determined by plaque assay as described above in BSC-1 cells.

### Statistical analysis

Statistical analyses were performed using either a two-tailed unpaired Student’s t-test (with Welch’s correction where variances were unequal), an ordinary one-way ANOVA with multiple comparisons, multiple *t*-tests, a Mantel-Cox test or a Mann-Whitney test as indicated in the figure legends. All analyses were performed with GraphPad Prism version 10. n.s. non-significant, * *P*<0.05, ** *P*<0.01, *** *P*<0.001

### Data availability

All data included in this study including supplementary material will be freely available. RNA-seq data will be uploaded at a relevant repository with appropriate dataset identifiers and can be made available upon request. OPXV nucleotide sequences cited in this study are available on NCBI GenBank: MPXV_CVR_S1 (ON808413), MPXV_CBM (GCA_964276875), MPXV_UK_P2 (MT903344), MPXV_UK_P3 (MT903345), VACV Western Reserve (AY243312), CPXV Brighton-Red (AF482758.2) and ECTV Moscow (AF012825).

## ACKNOWLEDGEMENTS

The authors thank K. Maringer, I. Dietrich, A. Zagrajek, S. Richardson and V. Sy (Pirbright Institute, UK) for assistance with the CL3 facility; M. Weekes and M. Potts (University of Cambridge, UK) for providing plasmid encoding MPXV OPG188; G.J. Towers (University College London, UK) for helpful discussions; and University of Glasgow CVR and UKHSA for providing MPXV isolates. This work was supported by grants BB/X0011356/1 and BB/V015265/1 from the UK Biotechnology and Biological Sciences Research Council to C.M.d.M; grants PID2022-136867NB-100 and RTI2021-128580OB-100 from the Spanish Ministry of Science and Innovation to B.H. and A.A., respectively; and grant HR23-00558 from La Caixa Foundation (Spain) to A.A. A.J.M.H. was supported by a PhD studentship from the Lorna and Yuti Chernajovsky Biomedical Trust.

## AUTHOR CONTRIBUTIONS

R.P.S. and C.M.d.M conceived the project. R.P.S., L.E., B.H., A.A. and C.M.d.M. designed and supervised the experiments. R.P.S., L.E., B.H, T.S.F., A.J.M.H., H.A., A.E., M.P., S.L., I.S., P.P, I.A., F.J.A., and A.R.U. performed the experiments and analysed data. B.C., G.L.S. and D.O.U. contributed key reagents and facilities. A.A., B.H. and C.M.d.M. acquired funding. R.P.S. and C.M.d.M wrote the original draft of the manuscript, with edits from B.H, A.A., G.L.S. and D.O.U. All authors revised and agreed to the final version of the manuscript.

## MATERIALS AND CORRESPONDENCE

Correspondence and requests for materials should be addressed to Carlos Maluquer de Motes.

