## Supplemental Data for "Attenuation of the 2022 global outbreak monkeypox virus relative to its clade IIb ancestor"

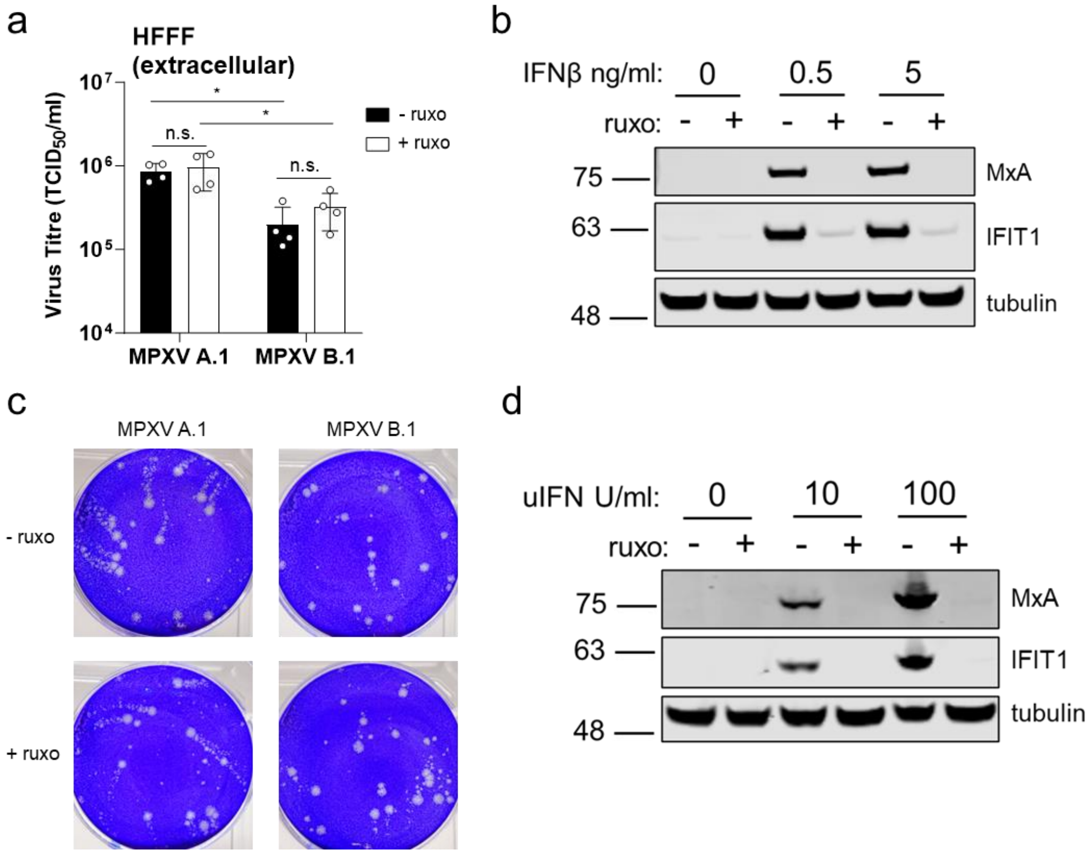


**Extended Data Figure 1: Effects of JAK1/2 inhibition on replication of MPXV Clade IIb lineages B.1 and A.1.**

**a**, MPXV A.1 and B.1 extracellular virus titre at 72 h in HFFF in the presence or absence of 2 μM ruxolitinib (ruxo). n = 4 per condition. Data are mean ± SD. Statistical analyses were performed using an ordinary one-way ANOVA with multiple comparisons. *$P < 0.05$, n.s. non-significant..

**b**, Immunoblot of HFFF cells treated for 16 h with the indicated concentrations of human IFNβ in the presence or absence of 2 μM ruxolitinib (ruxo).

**c**, Representative comet tail assay from BSC-40 cells infected with MPXV A.1 or B.1 for 96 h with a liquid overlay in the presence or absence of 1 μM ruxolitinib (ruxo).

**d**, Immunoblot of BSC-40 cells treated for 16 h with the indicated concentrations of universal IFN in the presence or absence of 1 μM ruxolitinib (ruxo).

All data presented are representative of at least 2 experimental repeats.

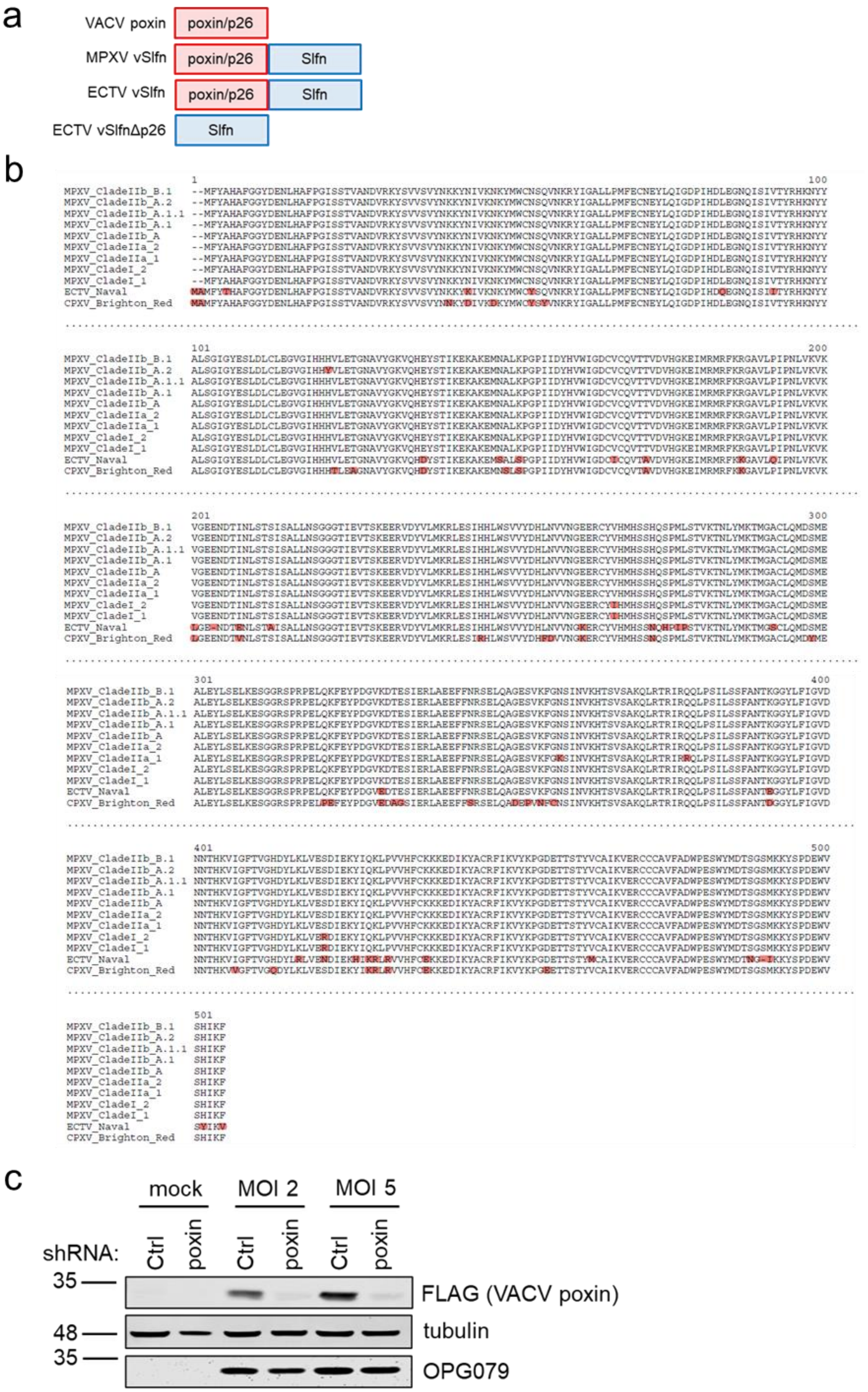
a
VACV poxin
poxin/p26
MPXV vSlfn
poxin/p26
Slfn
ECTV vSlfn
poxin/p26
Slfn
ECTV vSlfnΔp26
Slfn
b
MPXV_CladeIIb_B.1
MPXV_CladeIIb_A.2
MPXV_CladeIIb_A.1.1
MPXV_CladeIIb_A.1
MPXV_CladeIIb_A
MPXV_CladeIIa_2
MPXV_CladeIIa_1
MPXV_CladeI_2
MPXV_CladeI_1
ECTV_Naval
CPXV_Brighton_Red
c
mock
MOI 2
MOI 5
shRNA:
Ctrl
poxin
35
48
FLAG (VACV poxin)
tubulin
OPG079

**Extended Data Figure 2: MPXV utilises vSlfn to evade innate sensing.**

**a**, Schematic of the poxin/vSlfn constructs used in this study.

**b**, Amino acid alignment of OPXV vSlfn proteins with mismatches highlighted in red.

**c**, Immunoblot of THP-1-derived macrophages stably expressing either a control (Ctrl) or poxin-targeting shRNA, infected for 16 h with a VACV expressing FLAG-tagged B2 (poxin) at MOI 2 or 5.

Data in (c) is representative of 2 experimental repeats.

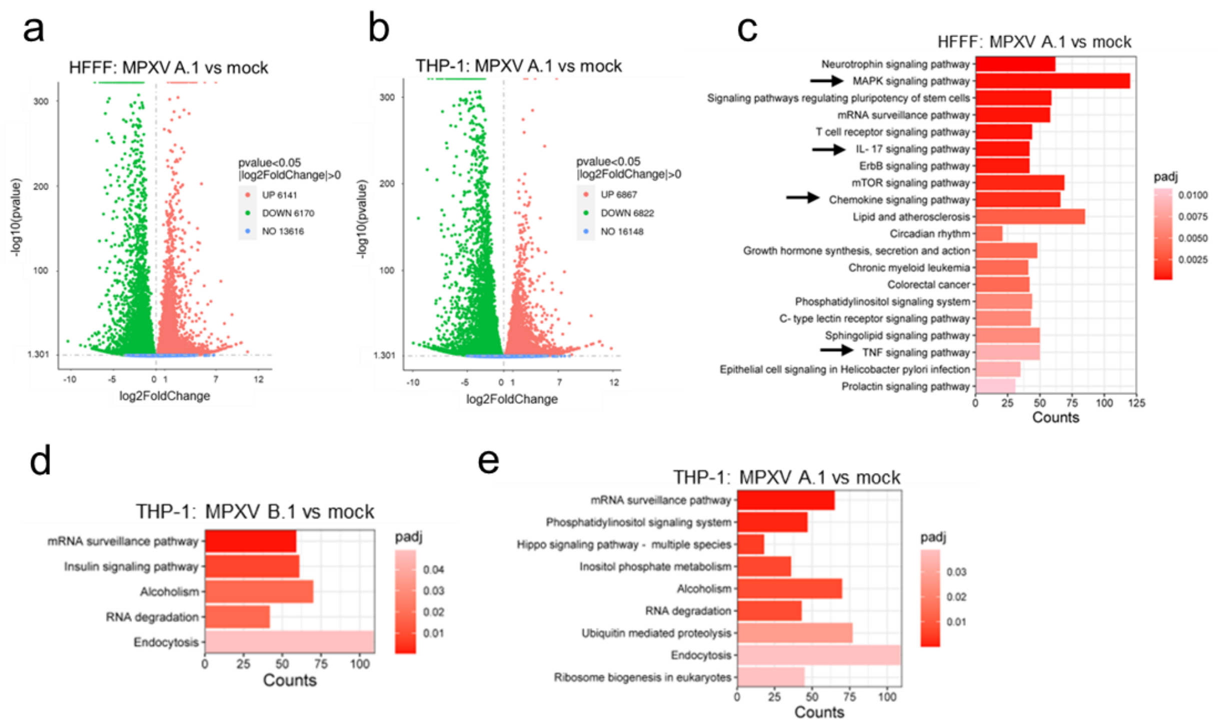


**Extended Data Figure 3: Transcriptional changes during MPXV infection.**

**a, b,** Significantly differentially expressed genes (log2FC>0.0) at 24 hpi by volcano plot analysis in HFFF (a) or THP-1-derived macrophages (b) infected with MPXV A.1 at MOI 5, compared with the mock. *P* value determined by DESeq2.

**c**, Top 20 significantly upregulated pathways (KEGG analysis) from RNAseq data of HFFF cells infected with MOI 5 MPXV A.1 for 24 h compared with mock-infected cells. *P* value determined by DESeq2. Pathways of interest are highlighted with a black arrow.

**d, e**, Significantly upregulated pathways (KEGG analysis) in THP-1-derived macrophages infected with MOI 5 MPXV B.1 (**d**) or A.1 (**e**) compared with mock-infected cells. *P* value determined by DESeq2. All data are from a single RNA-seq experiment.

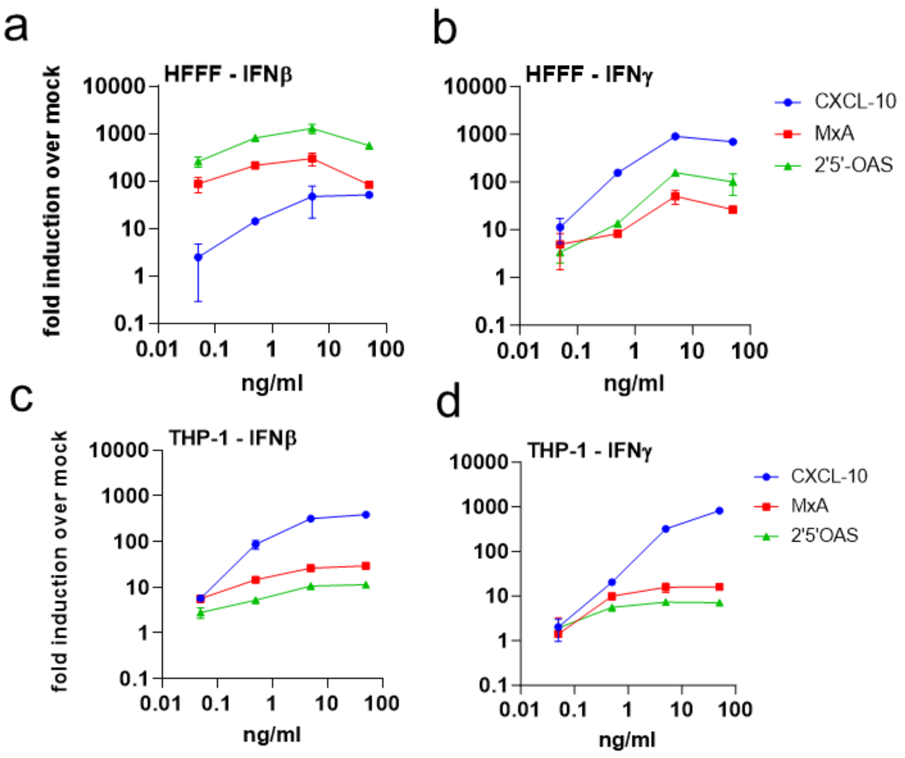


**Extended Data Figure 4: ISG responses in HFFF and THP-1-derived macrophages treated with type I or type II IFN.**

**a-d,** RT-qPCR for ISGs *CXCL-10*, *MxA* and *2'5'OAS* from HFFF (**a, b**) or THP-1-derived macrophages (**c, d**) treated for 16 h with the indicated doses of type I (IFNβ) (a, c) or type II (IFNγ) (b, d) IFN. All data are mean ± SD, n = 3 per condition.

Data presented are representative of 2 experimental repeats.

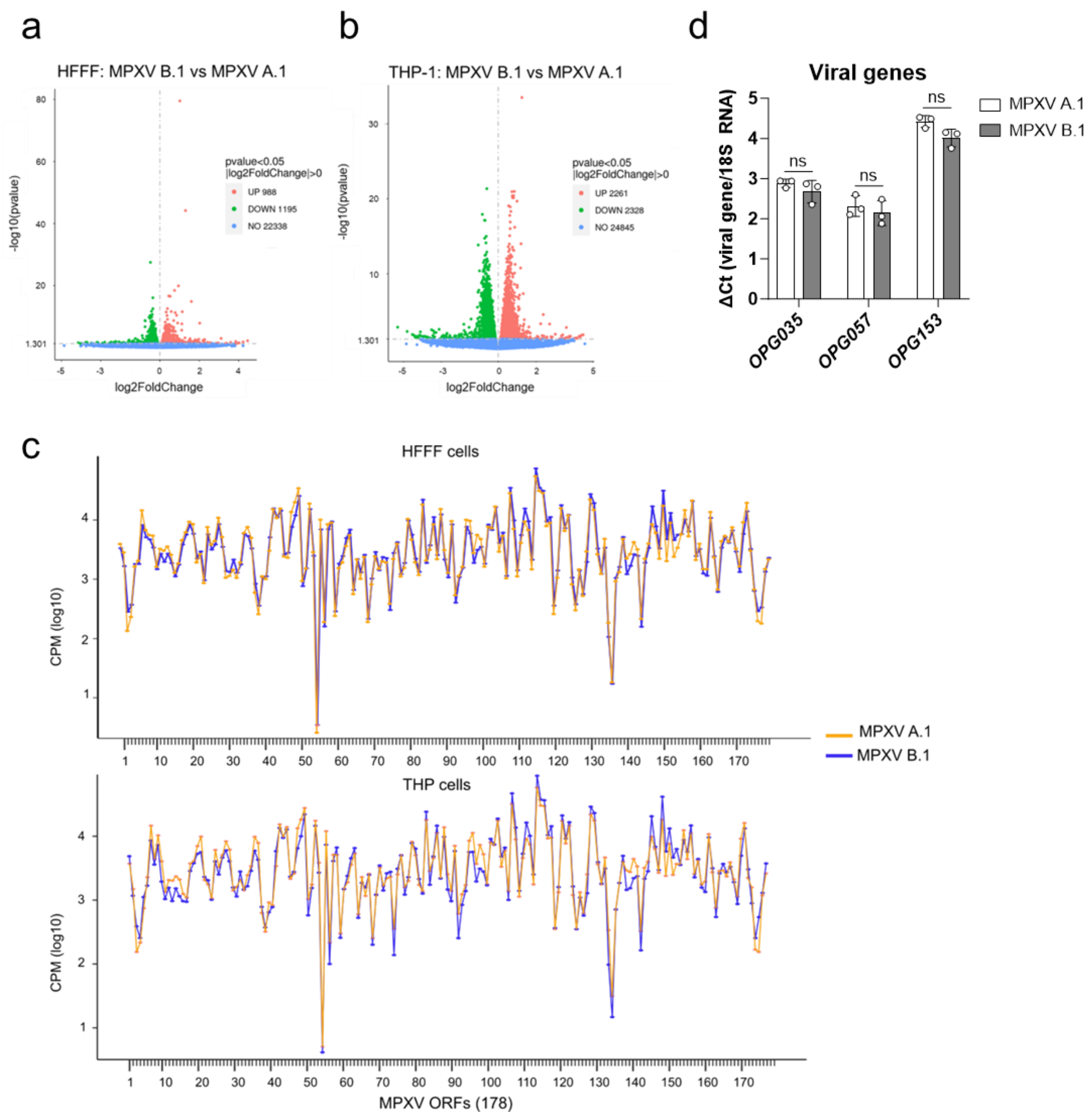


**Extended Data Figure 5: Differential innate immune control by MPXV lineages B.1 and A.1.**

**a, b**, Significantly differentially expressed genes (log2FC>0.0) in HFFF (**a**) and THP-1-derived macrophages (**b**) by volcano plot analysis at 24 hpi with MPXV B.1 at MOI 5 compared with MPXV A.1 infection. *P* value determined by DESeq2.

**c**, Counts per million (CPM) per MPXV ORF from RNA seq data.

**d,** Viral gene expression by RT-qPCR at 8 h post-infection from cells in Fig 4g measuring representative early (*OPG025)*, intermediate (OPG057*)* and late (OPG153*)* viral transcripts. Data are mean ± SD. n = 3 per condition. Statistical analyses were performed using an unpaired Student's t-test. ns, non-significant.

Data presented are from a single RNA-seq experiment, except (d) that is a representative experiment of 2 repeats.

**Supplementary Table 1**

**Single nucleotide differences between MPXV B.1 strain CVR_S1 (accession number ON808413) and MPXV A.1 strain UK_P3 (accession number MT903345)**

| Position | A.1 nt | B.1 nt | A. 1 aa | B.1 aa | OPG gene | VACV COP gene | Description |
|---|---|---|---|---|---|---|---|
| 1284 | G | A | S | L | OPG001 | C23L | secreted chemokine binding protein |
| 2613 | G | A | S | F | OPG002 | C22L | secreted TNFα receptor like protein |
| 3133 | G | A | non-coding | | OPG003 | C19L | ankyrin repeat protein |
| 3544 | G | A | non-coding | | OPG003 | C19L | ankyrin repeat protein |
| 3840 | C | T | D | N | OPG003 | C19L | ankyrin repeat protein |
| 7793 | C | T | non-coding | | OPG019 | C11R | EGF like domain protein |
| 14022 | G | T | A | D | OPG025 | C9L | ankyrin repeat protein |
| 15450 | G | A | intergenic | | | | |
| 21745 | G | A | non-coding | | OPG037 | M1L | ankyrin-like protein |
| 30389 | G | A | non-coding | | OPG047 | F3L | kelch-like protein |
| 31075 | G | A | R | C | OPG047 | F3L | kelch-like protein |
| 34481 | G | A | P | S | OPG053 | F9L | IMV membrane protein |
| 37224 | G | A | non-coding | | OPG056 | F12L | EEV maturation protein |
| 38382 | G | A | non-coding | | OPG056 | F12L | EEV maturation protein |
| 38684 | C | T | E | K | OPG056 | F12L | EEV maturation protein |
| 39161 | C | T | E | K | OPG057 | F13L | Palmytilated EEV membrane protein |
| 52907 | G | A | non-coding | | OPG071 | E9L | DNA polymerase |
| 54139 | G | A | L | F | OPG071 | E9L | DNA polymerase |
| 54657 | G | A | D | N | OPG072 | E10R | sulfhydryl oxidase |
| 64319 | G | A | non-coding | | OPG083 | I7L | virion core cysteine protease |
| 73088 | C | T | S | L | OPG093 | G8R | VLTF-1 late transcription factor |
| 73261 | G | A | D | N | OPG093 | G8R | VLTF-1 late transcription factor |
| 74227 | G | A | M | I | OPG094 | G9R | entry/fusion complex component |
| 77405 | G | A | E | K | OPG098 | L4R | ss/dsDNA binding protein |
| 81297 | G | A | non-coding | | OPG105 | J6R | RNA polymerase subunit (RPO147) |
| 82395 | C | T | non-coding | | OPG105 | J6R | RNA polymerase subunit (RPO147) |
| 82473 | G | A | non-coding | | OPG105 | J6R | RNA polymerase subunit (RPO147) |
| 84609 | C | T | non-coding | | OPG105 | J6R | RNA polymerase subunit (RPO147) |
| 95056 | G | A | non-coding | | OPG115 | D3R | virion core protein |
| 124152 | G | A | E | K | OPG145 | A18R | DNA helicase |
| 124696 | G | A | R | Q | OPG145 | A18R | DNA helicase |
| 128720 | C | T | S | L | OPG150 | A23R | intermediate transcription factor VITF-3 |
| 150494 | C | T | H | Y | OPG176 | A46R | IL-1/TLR signaling inhibitor |
| 151486 | A | C | intergenic | | | | |
| 155820 | G | A | intergenic | | | | |
| 162356 | C | T | non-coding | | OPG188 | B2R | viral schlafen |
| 170287 | G | A | non-coding | | OPG198 | B12R | Ser/Thr Kinase |
| 178204 | G | A | intergenic | | | | |
| 181997 | G | A | D | N | OPG210 | B21R | surface glycoprotein |
| 183536 | C | T | P | S | OPG210 | B21R | surface glycoprotein |

| | | | | | | | |
|---|---|---|---|---|---|---|---|
| 186696 | G | A | M | I | OPG210 | B21R | surface glycoprotein |
| 193409 | G | A | D | N | OPG003 | C19L | ankyrin repeat protein |
| 193705 | C | T | non-coding | | OPG003 | C19L | ankyrin repeat protein |
| 194116 | C | T | non-coding | | OPG003 | C19L | ankyrin repeat protein |
| 194636 | C | T | S | F | OPG002 | C22L | secreted TNFα receptor like protein |
| 195965 | C | T | S | L | OPG001 | C23L | secreted chemokine binding protein |
